# Inhibition of the metalloprotease ADAM19 as a Novel Senomorphic Strategy to Ameliorate gut permeability and senescence markers by modulating senescence-associated secretory phenotype (SASP)

**DOI:** 10.1101/2024.12.16.628718

**Authors:** Sudipta Bar, Tyler A.U. Hilsabeck, Baline Pattavina, Jose Alberto Lopez-Dominguez, Nathan Basisty, Joanna Bons, Mark Watson, Birgit Schilling, Judith Campisi, Pankaj Kapahi, Amit Sharma

**Author notes:** Equal contributing co-corresponding authors.

## Abstract

Accumulation of DNA damage can accelerate aging through cellular senescence. Previously, we established a *Drosophila* model to investigate the effects of radiation-induced DNA damage on the intestine. In this model, we examined irradiation-responsive senescence in the fly intestine. Through an unbiased genome-wide association study (GWAS) utilizing 156 strains from the Drosophila Genetic Reference Panel (DGRP), we identified meltrin β (the drosophila orthologue of mammalian ADAM19) as a potential modulator of the senescence-associated secretory phenotype (SASP). Knockdown of *meltrin β* resulted in reduced gut permeability, DNA damage, and expression of the senescence marker β-galactosidase (SA-β-gal) in the fly gut following irradiation.

Additionally, inhibition of ADAM19 in mice using batimastat-94 reduced gut permeability and inflammation in the gut. Our findings extend to human primary fibroblasts, where ADAM19 knockdown or pharmacological inhibition decreased expression of specific SASP factors and SA-β-gal. Furthermore, proteomics analysis of the secretory factor of senescent cells revealed a significant decrease in SASP factors associated with the ADAM19 cleavage site. These data suggest that ADAM19 inhibition could represent a novel senomorphic strategy.

## Introduction

One of the most prevailing hypotheses for aging is that it results from accumulated unresolved DNA damage (approximately 2 x 10^5^ lesions per day)[1] due to a progressive decline in the DNA repair processes, culminating in age-related frailty and tissue dysfunction[2–8]. One consequence of unresolved and erroneous DNA repair is a state of replicative arrest, termed cellular senescence[9]. Although this response can prevent malignant transformation, it can also contribute to aging because senescent cells, in addition to contributing to tissue dysfunction, secrete pro-inflammatory SASP and create a pro-inflammatory microenvironment. Understanding the mechanisms by which senescent cells contribute to a decline in tissue function is critical for developing novel therapeutic interventions for age-associated disorders. Several groups have developed senolytics to kill senescent cells selectively[10–12]. As the SASP secreted by senescent cells is known to drive inflammaging or age-related sterile inflammation[13–15] thus, in addition to removing senescent cells, inhibiting the SASP is also a viable strategy to limit the detrimental effects of senescence in aging.

We have recently published the *Drosophila melanogaster* model to study the molecular effects of radiation-induced damage in the intestine flies[16]. In addition, we performed an unbiased GWAS screen (using 156 strains from the *Drosophila* Genetic Reference Panel) to search for natural genetic variants that regulate radiation-induced gut permeability in adult *D. melanogaster*[17]. We have previously reported the role of an RNA-binding protein, *Musashi* (*msi*), in regulating the proliferation of intestinal stem cells following the damage caused by ionizing radiation in the gut of adult flies[16].

In this study, we report another candidate gene *meltrin β* from our genetic screen with ∼156 DGRP strains with naturally occurring variations to identify novel modulators of the tissue repair pathways following DNA damage by screening for gut permeability after irradiation. It belongs to a class of type I transmembrane glycoproteins called <u>A D</u>isintegrin <u>a</u>nd <u>M</u>etalloproteases (ADAMs). ADAM(s) are highly conserved in several species, and their primary function is domain shedding[18] [19]. Members of this class have a wide range of regulatory functions. They can proteolytically release several cytokines, growth factors (TNF-α, TGF-β), receptors (IL-1R-II), adhesion molecules, and enzymes from the plasma membrane[20–25]. ADAMs, hasve emerged as an essential player in cancer progression and metastasis[26,27]. where its inhibition is suggested to have a potential therapeutic value.

Here, we demonstrate that knocking down *ADAM19/meltrin β* or its inhibition using a small molecule, Batimastat (BB-94), reduces gut permeability and some senescence-associated phenotype in irradiated flies. We also observed an increase in the expression of β Galactosidase and Upd3 in the cells gut of irradiated flies with meltrin knockdown. Interestingly, inhibition of ADAM19 by Batimastat (BB-94) also reduced intestinal permeability in doxorubicin-treated mice. We also observed that BB-94 treatment reduced proinflammatory SASP in the colon of doxorubicin-treated mice. Additionally, our proteomics analysis of the secretome following ADAM19 knockdown (or pharmacological inhibition) in irradiated human primary fibroblasts reduced several SASP factors and Senescence-Associated β-Gal (SA-β-Gal) expression. Finally, bioinformatic analysis of the proteomics data confirmed conserved ADAM19 cleavage sites in several SASP factors significantly reduced when ADAM19 was knocked down in senescent cells.

In summary, we used flies as a rapid platform to discover new genes (e.g., *meltrin β*) that modulated the SASP or senescence (in mammals) and showed the potential of reducing SASP expression instead of eliminating senescent cells by ADAM19 inhibition as a novel senomorphic strategy.

## Results

### Increase in the expression of persistent DNA damage and β galactosidase expression in the gut of flies

Several lines of evidence have demonstrated beyond doubt that DNA damage accumulates with age[28]. Increased reactive oxygen species production and a decline in DNA repair capacity have been implicated but completely understood and has been the focus of research[28]. To gain insight into underlying mechanisms at both molecular and cellular levels, we leveraged *Drosophila melanogaster* to investigate the molecular effects of exposure to ionizing radiation on intestinal permeability in flies in our previous publication[16].

The 5-day-old adult female w^1118^ flies were exposed to ionizing radiation (100 Gy) that is known to induce DNA damage. The cells undergoing DNA double-strand breaks are known to activate the DNA damage response (DDR)[29]. This repair response can be reliably measured by quantifying the histone variant H2AX phosphorylation, producing γH2AX[29]. We performed the immunofluorescence assay in the dissected gut seven days after irradiation and observed a significant increase in cells expressing γH2AX. We also observed a concomitant increase in β-galactosidase expression by co-immunostaining the fly gut following dissection seven days after irradiation (**Figure 1A, Ai and Aii**). Interestingly, we also observed a ∼7-fold increase in the expression of β-galactosidase and a higher percentage of cells expressing γH2AX (42% of cells were γH2AX-positive compared to 0.4% in the control) in the gut of w1118 flies in an age-dependent manner, as shown by our immunofluorescence assay in the gut of twenty-day-old flies compared to seven-day-old control flies. (**Figure 1B, B-i and B-ii**).

**Figure 1:**
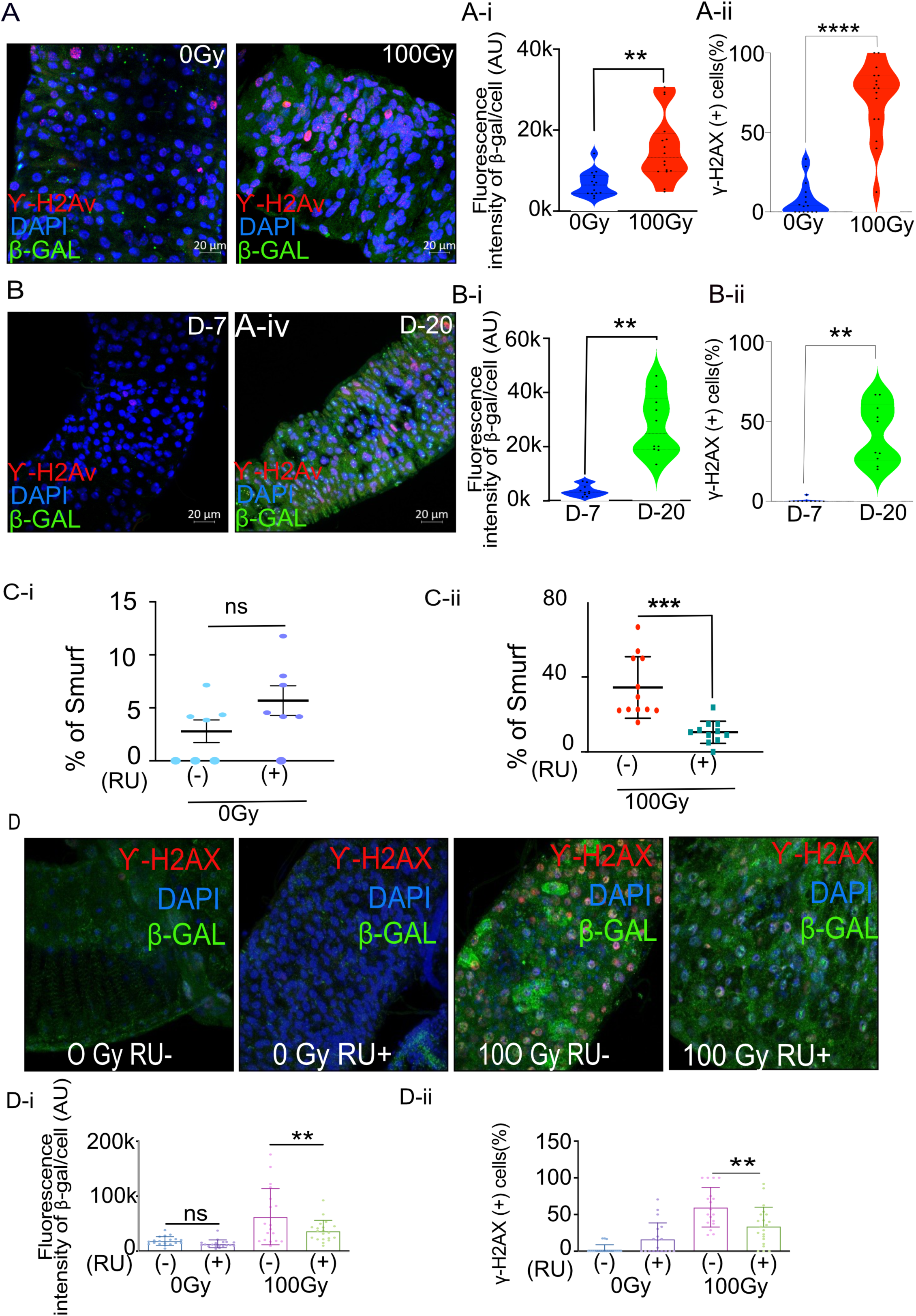
Increase in senescence makers in irradiated and aged flies and *meltrin β* knockdown in the enterocytes reduces intestinal permeability and senescence makers in flies following irradiation. (A) Representative immunohistochemistry images of the gut show increased SA-β gal and γ-H2Ax positive cells in X-ray irradiated flies (100Gy) compared to control flies (0Gy). Fluorescence intensity was measured using ImageJ; Scale bar = 20 μm. **(Ai).** Quantification of fluorescence intensity of SA-β gal (arbitrary units, A.U.) between 0 Gy and 100 Gy. Mean fluorescence intensity was measured using ImageJ and averaged across the gut of 15-16 flies. Data represent mean ± SEM for each group. **(Aii)** Quantification of γ-H2AX positive cells expressed as a percentage of total cells in both control and 100 Gy X-ray exposed groups. Data are presented as mean ± SEM from the gut of 15-16 flies. Exposure to 100 Gy of X-ray radiation significantly increased the number of γ-H2AX positive cells compared to the control group, indicating elevated DNA double-strand breaks in response to radiation. **(B)** Representative immunohistochemistry images of the gut show increased SA-β gal and γ-H2Ax positive cells in X-ray irradiated flies (20 days) compared to control flies (7 days). Fluorescence intensity was measured using ImageJ; Scale bar = 20 μm. **(Bi)** Quantification of fluorescence intensity of SA-β gal (arbitrary units, A.U.) between 7-day and 20-day-old flies. Mean fluorescence intensity was measured using ImageJ and averaged across the gut of 13-16 flies. Data represent mean ± SEM for each group. **(Bii)** Quantification of γ-H2AX positive cells expressed as a percentage of total cells in both 7-day-old and 20-day-old flies. Data are presented as mean ± SEM from the gut of 13-16 flies. The gut of 20 days old flies significantly increased the number of γ-H2AX positive cells compared to the 7-day-old, indicating elevated DNA double-strand breaks in response to radiation. **(Ci and Cii)** Smurf assay for assessing gut permeability was performed with *5966-GS>UAS-Meltrin RNAi* flies on day ten after irradiation. Control 100Gy R (-) was maintained on 1.5 YE food without RU486, whereas *meltrin β* expression was knocked down in ECs in 100Gy R (+). Results plotted as the mean proportion of smurf to non-Smurf flies of 4 vials in each group with 25 flies in each vial. Error bars indicate SEM. **(D)** Immunohistochemistry images of gut showing an increase in SA-β gal and γ-H2Ax positive cells in non-irradiated and X-Ray irradiated in control (RU-) flies and *meltrin β* RNAi and flies (RU+). **(D-i and D-ii)** Quantification of SA-β gal intensity and γ-H2Ax positive cells in irradiated-non irradiated flies in the gut of RU- and RU+ flies. **(A-i, A-ii, B-i, B-ii, C, C-i, D-i and D-ii)** Error bars indicate SEM. The significance level was determined by ANOVA or Student’s t-test, denoted by asterisks: * for p < 0.05, ** for p < 0.01, *** for p < 0.001, and **** for p < 0.0001.

### Meltrin β knockdown in the gut reduces gut permeability, DNA damage, and β galactosidase expression

Our recent publication reported a GWAS analysis and identified several genes associated with increased intestinal permeability following radiation exposure. Among the candidates identified was meltrin[16]. Interestingly, the q-RTPCR indicated a significant decrease in the expression of meltrin in the gut of flies ten days following the exposure to ionizing radiation (100Gy) (Supplementary Fig.1A).

To investigate if *meltrin β* may influence gut permeability, DNA damage, and senescence markers after irradiation, we used the Gal4-UAS system to knock down *meltrin β* expression in enterocytes (ECs)[30]. RNAi targeting meltrin β was expressed using the drug RU486 inducible (RU+), EC-specific 5966-GS driver in 5-day-old adult flies, with vehicle-treated uninduced control flies (RU-) used for comparison. Meltrin β knockdown was confirmed by quantitative real-time PCR (Supplementary Fig. 1B). The Smurf assay was performed to determine the effect of *meltrin β* as previously described[31]. Briefly, this assay involves feeding flies a blue dye, and in a few hours, if the intestine is damaged and permeable, the dye diffuses to the hemolymph, and the whole fly appears blue. Knocking down meltrin β in ECs (RU+) in non-irradiated control flies showed no significant changes in gut permeability upon meltrin β knockdown (Fig. 1C-i). In contrast, followed by irradiation (100 Gy + RU) resulted in a reduction in intestinal permeability, as we observed in a 2-fold lower percentage of smurf flies compared to irradiated control flies without RU486 (-RU) exposed to 100 Gy. **(Fig. 1C-ii)**.

This metalloprotease inhibitor batimastat (BB-94) can inhibit several memebers of ADAM family[32,33]. Hence, to phenocopy the effects of its knockdown of ADAM 19 (meltrin) on intestinal permeability after irradiation, we treated w1118 flies with 50 μM BB-94 in the food. We observed significantly reduced intestinal permeability in w1118 female flies 14 days after irradiation (Supplementary Fig. 1C). As exposure to ionizing radiation is known to cause DNA damage, we tested the effect of knocking down meltrin β in ECs on the expression γH2AX and β-galactosidase. As expected, our results show a significant increase in the mean fluorescence intensity of β-galactosidase expression and percentage of cells expressing γH2AX in control flies upon exposure to 100Gy of ionizing radiation RU486 (RU-), whereas knocking down meltrin β in ECs (RU+), followed by irradiation (100Gy), significantly reduced β-galactosidase expression, and the percentage of cells expressing γH2AX compared to the no RU486 (RU-) control. At the same time, non-irradiated flies did not show a change in the expression of either γH2AX or β-galactosidase in their gut with or without meltrin β in ECs (**Fig. 1D, 1D-i, and 1D-ii**).

As we reported in our previous publication, damage caused by exposure to ionizing radiation significantly decreases the gut size in adult W^1118^ female flies. Presumably, it is because of an increase in the apoptotic clearance of at least three days [16]. Using the same approach of dual acridine orange/ethidium bromide (AO/EB) fluorescent staining to quantify apoptosis-associated changes of cell membranes during the process of apoptosis [16]our results demonstrate a higher number of AO/EB positive cells in the gut of flies upon meltrin knockdown (RU+) on day five after irradiation compared to the control (RU-) (Supplementary Fig. 1D). In contrast, apoptotic cells decreased significantly at day 10 (Supplementary Fig. 1E).

Exposure to ionizing radiation is known to increase intestinal permeability and reduce survival in w^1118^ flies[16]. As ECs-specific knockdown of meltrin reduced intestinal permeability, we investigated if gut-specific knockdown of meltrin may improve survival and observed no effect on the mean or median lifespan of irradiated and non-irradiated flies (Supplementary Fig. 1F).

### *Meltrin β* /ADAM19 inhibition in the gut reduces gut permeability in mice model

As mentioned earlier, *meltrin β* is a highly conserved gene, and its mammalian homolog is ADAM19[34,35]. Interestingly, ADAM19 is known to be elevated in intestinal cells in inflammatory bowel disease (IBD) patients[36]. We wondered if ADAM19/*meltrin β* may also regulate intestinal permeability in mammals, especially in response to the damage caused by genotoxic stress. To test this, we used the well-established chemotherapy model. Doxorubicin(doxo) is a well-known chemotherapy agent that induces DNA damage by inhibiting topoisomerase II[37]. However, as a chemotherapy drug, a side effect of doxo is increased intestinal permeability [38,39]. As ADAM19 knockout mice die soon after birth due to cardiac defects[40], we used a BB-94 to pharmacologically inhibit its function, testing our hypothesis.

We treated four-month-old C57BL6 mice with or without doxo as a single intraperitoneal (*i.p.)* injection (doxo) (10 mg/kg) or similarly treated with vehicle controls (Ctrl) described previously[41]. BB-94 was administered with *i.p* injection (5 or 20 mg/Kg, once a week for six weeks) **(Fig. 2A)**. Intestinal permeability was measured by performing ELISA to detect fecal albumin, frequently used as a proxy for intestinal permeability[42]. We performed an ELISA test on the fecal material collected from the mice six weeks after doxo treatment and detected an increase in the amount of albumin detected in the fecal matter compared to vehicle-treated control mice. However, mice treated with 5 and 20 mg/Kg BB-94 had significantly lower albumin in the fecal matter compared to doxo-treated mice **(Fig. 2B)**. There was no statistical difference between mice treated with 5 or 20 mg/Kg BB94 and the amount of albumin detected in the fecal matter collected from mice treated with either **(Fig. 2B)**.

**Figure 2:**
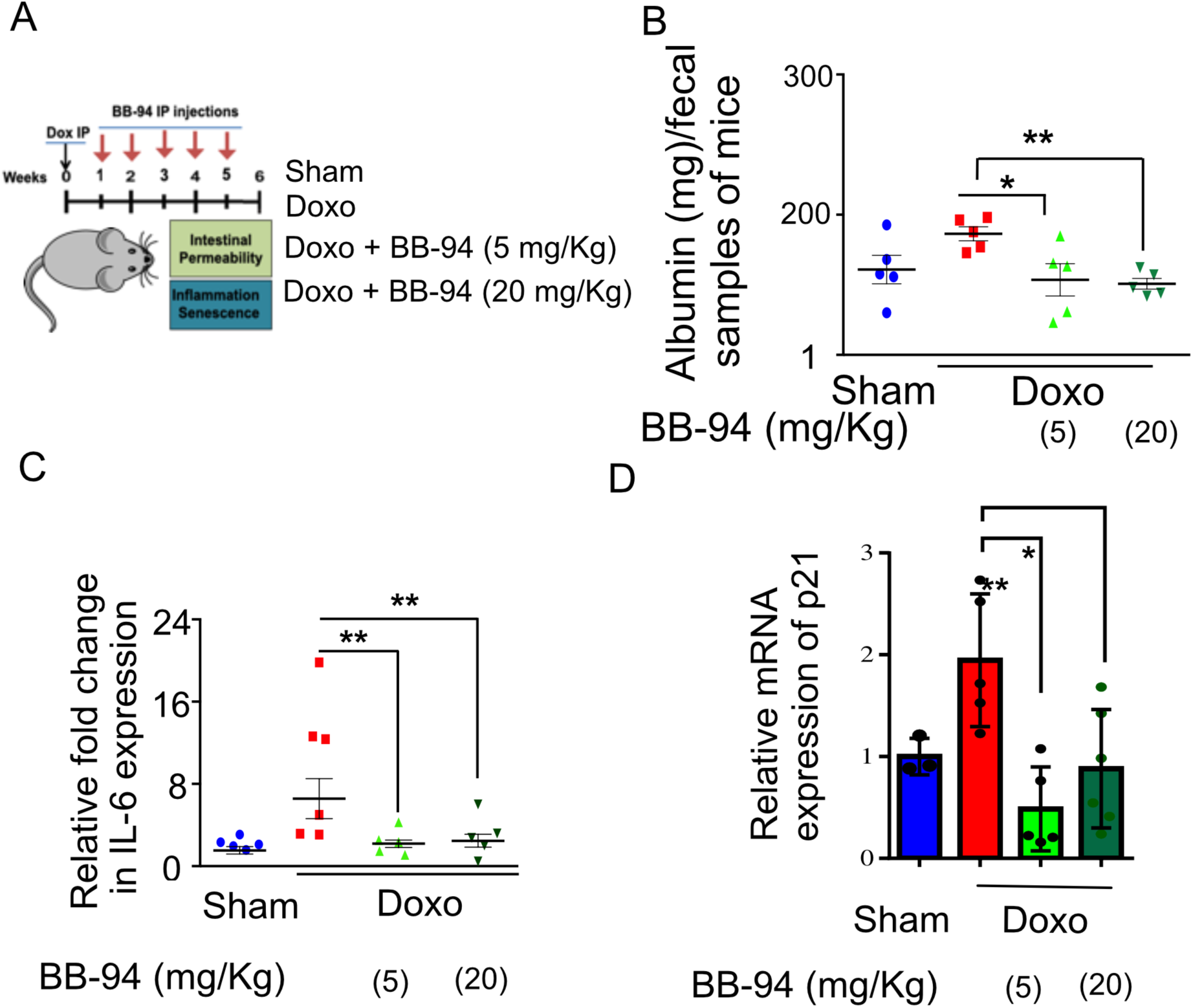
ADAM19 inhibition by BB-94 treatment reduces intestinal permeability and inflammation in mice caused by Doxorubicin treatment. **(A)** Experimental design to determine the effect of ADAM19 inhibition in a mouse model. Mice received a single dose, i.p Dox (10 mg/Kg) or vehicle (SHAM Ctrl) at the start of the experiment. Followed by a subset of mice receiving Bataimastat BB-94 doses of 5 mg/Kg and 20 mg/Kg once a week, every other week till week 5. Intestinal permeability and inflammation were measured on week six in the colon of dissected mice. **(B)** Albumin levels were measured using an ELISA assay in early morning fecal samples from each group. The graph presents albumin levels as mean ± SEM, expressed in mg per 100 mg/ml of fecal material, from (n=6) across. Significant differences in albumin levels between groups (e.g., Sham vs doxo, doxo vs BB-94 treated mice at 5 mg/kg and 20 mg/kg, etc.) were assessed using ANOVA, with p-values indicated as *p < 0.05, **p < 0.01, ***p < 0.001. Elevated albumin levels in feces may indicate increased intestinal permeability or gut damage, reflecting potential gastrointestinal dysfunction in doxo-treated mice, while treatment with BB-94 significantly reduced it. **(C)** Real-time PCR was performed with RNA prepared from dissected colon tissue of mice treated as described above. The results are presented as mean relative fold change in the gene expression of IL-6 and normalized to the housekeeping gene Actin. **(D)** Mean relative fold change in the gene expression of p21 and normalized to the housekeeping gene Actin **(B-D)** Error bars indicate SEM. The significance level was determined by ANOVA and denoted by asterisks: * for p < 0.05, ** for p < 0.01, *** for p < 0.001, and **** for p < 0.0001.

Doxorubicin treatment is known to increase the inflammatory response in various tissues in mice[43,44] we investigated the effect of BB-94 on the inflammation in the gut. Our real-time PCR results in the colon tissue of the mice confirmed an increase in interleukin-6 (IL-6) expression upon doxorubicin treatment. However, mice that received either concentration BB-94 (5 or 20 mg/Kg) had significantly reduced expression of IL-6. **(Fig. 2C)**. Interestingly, IL-6 is the mammalian homolog of Upd3, which also increases in the expression in the gut of flies upon exposure to genotoxic stress. A similar trend was observed in TNFα expression, although it was not statistically significant (p ≤ 0.05) due to the small sample size (**Supplementary Fig. 2A**).

The CDK inhibitor p21 regulates p53-mediated growth arrest following DNA damage[45]. Interestingly, our qRT-PCR in doxorubicin-treated mice showed a trend of increased cell cycle checkpoint maker p21 expression in the colon, which was significantly reduced by both doses (5 mg/kg and 10 mg/kg) of BB-94 (**Fig. 2D**).

### ADAM19 modulates cellular response to genotoxic stress

In response to genotoxic stressors like doxorubicin or irradiation, mammalian cells that accumulate persistent DNA damage can permanently escape the cell cycle by a phenomenon described as cellular senescence[46]. Senescent cells are also known to secrete a complex mix of pro-inflammatory cytokines, chemokines, growth factors, and matrix-modifying proteins collectively called the SASP[47].

ADAM19 inhibition reduced intestinal permeability and inflammation by reducing the senescence phenotype and proinflammatory upd3 and β-galactosidase in flies and il6 and p21 expression in mice colon. We then reasoned if ADAM19 may regulate senescence phenotype. To this end, we tested the effect of ADAM19 knockdown or inhibition in human lung fibroblasts IMR-90 doxorubicin (250 nMol, a well-established model of cellular senescence. We observed a two-fold increase in the expression of ADAM19 in doxo-treated S cells. Using shRNA, we observed a significant decrease in its expression (Supplementary Fig 3A).

The quantitative real-time PCR indicates a significant increase in the expression of pro-inflammatory SASP *il1α, il6, and il8*, whereas ADAM19 knockdown or its inhibition by BB94 significantly reduced the expression of these SASP factors, with on effect of control shRNA **(Fig. 3A and Supplementary Fig. 3B and C)**.

**Figure 3.**
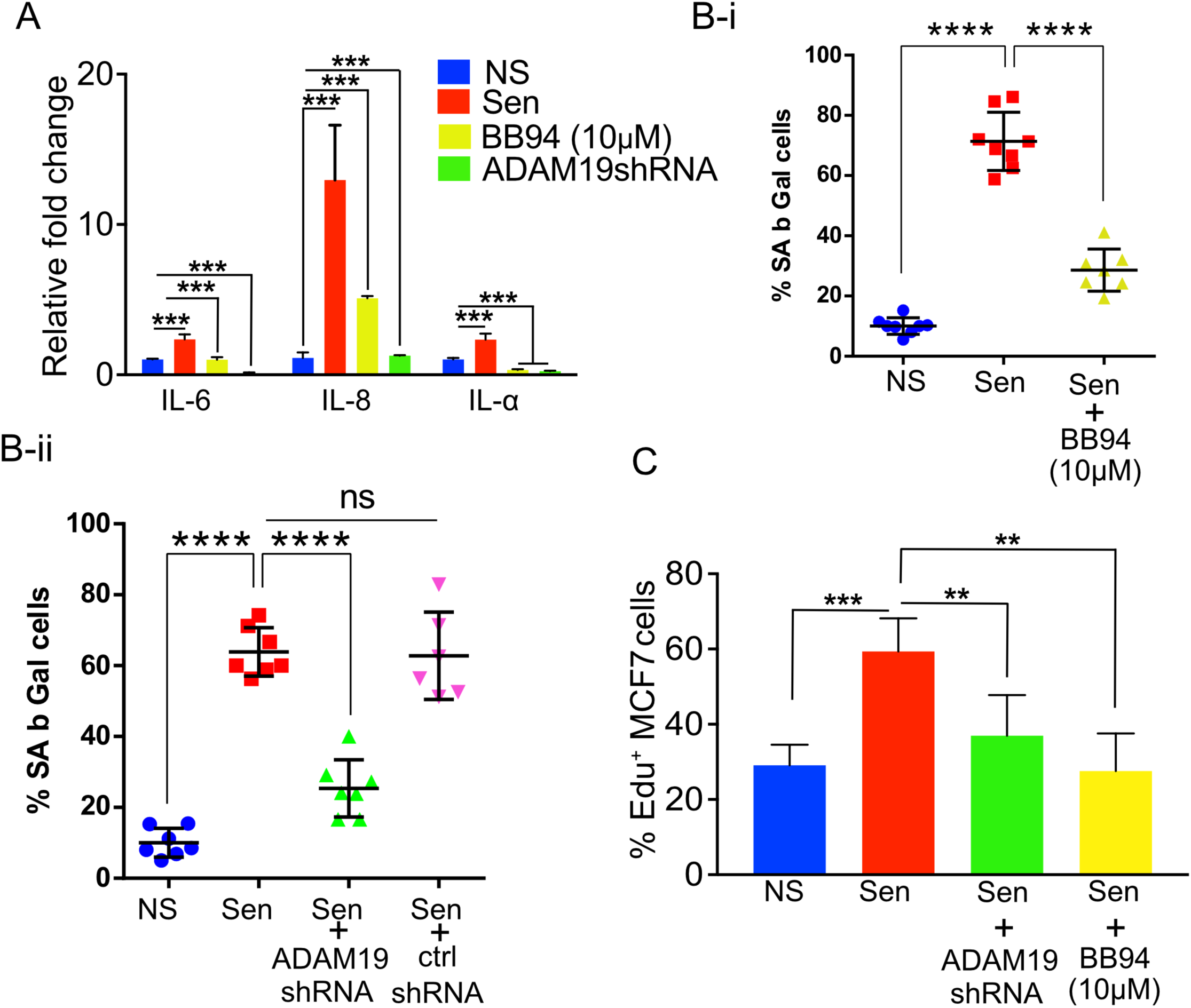
ADAM19 knockdown and pharmacological inhibition reduced senescence in human fibroblast cells caused by irradiation. (A) The mean fold change in the expression levels of senescence-associated secretory phenotype (SASP) cytokines IL-6, IL-8, and IL-1α was measured by real-time PCR (qPCR) in IMR-90 cells ten days after exposure to 100 Gy X-ray. The expression levels were quantified relative to untreated control samples and normalized to the housekeeping gene actin (ACTB) to control for differences in input RNA and reverse transcription efficiency. Data are presented as mean ± SEM from three independent experiments. The results represent the relative fold change in gene expression compared to control. Increased expression of SASP cytokines, such as IL-6, IL-8, and IL-1α, indicates a senescence response and may reflect the inflammatory environment of irradiation-induced cellular damage, which was significantly reduced in cells with ADAM19 knockdown or inhibition following treatment with BB-94. (B-i) IMR-90 cells were subjected to SA β-Gal staining ten days after irradiation (10Gy), following 10 µM BB-94 treatment. (B-ii) IMR-90 cells were subjected to SA β-Gal staining ten days after irradiation (10Gy), following transduction with either ADAM19 or scrambled shRNA. The percentage of SA β-Gal positive cells was quantified and presented as mean ± SEM, with three independent experiments with two technical replicates per group. (C) MCF-7 cells were treated with conditioned media collected from IMR-90 cells ten days after irradiation with or without ADAM19 shRNA or with or without Batimastat (10 μM). Results plotted as mean % EdU positive cells. **(A-C)** Error bars indicate SEM. The significance level was determined by ANOVA and denoted by asterisks: * for p < 0.05, ** for p < 0.01, *** for p < 0.001, and **** for p < 0.0001. In A, B-I, B-ii and C “NS” represent non senescence and ‘Sen’ represent senescence.

The lysosomal β-galactosidase activity in senescent cells is well documented and often quantified to measure senescence. We observed an increase 6-fold in the percentage of β-galactosidase-expressing cells in senescent cells (Sen) ten days after doxorubicin treatment. As expected, we observed a significant increase in the expression of SA β-gal in the proportion of cells treated with doxo (10 days after doxo treatment) compared to non-senescence (NS) control cells. While knocking down ADAM19 significantly decreased the cell activity of SA β-gal (from 64% to about 25%). Control shRNA did not affect the β-galactosidase-expressing cells **(Fig. 3B-i)**. Similarly, BB-94 treatment (10μM) significantly lowered the percentage of cells expressing β-galactosidase compared to vehicle-treated Sen cells (from about 70% to about 30%) **(Fig. 3B-ii)**.

The expression of Lamin B1 is known to be diminished in senescent cells **(Supplemental Fig. 3D)**[48]. However, ADAM19 knockdown or inhibition did not restore. Also, the loss of HMGB1 from the nucleus of the senescent cells is also used to characterize senescence, and we did not observe any improvement in nuclear localization of HMGB1 in senescent cells with ADAM19 knockdown **(Supplemental Fig. 3E)**[49].

The SASP, secreted by senescent cells, is often attributed to causing systemic inflammation[50]. Chronic exposure to SASP is known to drive paracrine senescence[51]. However, paradoxically, exposure to proinflammatory SASP can create a pro-carcinogenic microenvironment and the development of aging-associated cancers[52]. Previously, cancer cell lines cultured with SASP collected from senescent cells were shown to proliferate at a faster rate. Our results agree with the published literature, as shown by the EdU staining of MCF7 cells grown with culture media (CM) collected from senescent cells proliferated two times faster than CM collected from non-senescent cells. However, the proliferation of MCF7 cells was significantly reduced when grown in CM media collected from senescent cells with ADAM19 knockdown or inhibition as measured by EdU staining performed two days after treatment with the CM collected from senescent or non-senescent IMR-90 cells treated with or without ADAM19 knockdown or inhibition with Batimastat (10 μM), (**Fig. 3C**).

### ADAM19 in modulating the SASP secretion from senescent cells

ADAM19 is known to be a sheddase and is primarily shown to be involved in ectodomain shedding[20]. For a comprehensive understanding of the effect of ADAM19 on the SASP composition, we performed proteomic analysis (data-independent acquisitions, DIA[53,54] of the CM (conditioned media) collected with senescent or non-senescent IMR-90 cells following ADAM19 knockdown or inhibition with or without Batimastat (**Fig. 4A**). We quantified a total of 472 proteins (**Supplementary spread sheet 1**). We determined the overlap in proteins that contain the *meltrin β*/ADAM 19 cleavage site and fold changes having corrected q-values of 0.05 or lower in comparisons between ADAM19 knockdown vs. control (a19 vs. ctrl: 82 proteins) and ADAM19 -irradiated vs control irradiated (a19-IR vs. IR: 30 proteins) (**Fig. 4B-I and B-ii**). Using the hyperfunction in R to calculate the probability of observing an overlap of 12 proteins between these two groups, we found that this overlap was unlikely to have occurred by chance, given the total number of proteins quantified (p-value of 3.68e-4). To identify a common senescence-specific ADAM19 binding motif among the overlapping proteins with significant fold changes, we performed motif analysis (https://myhits.sib.swiss/), which found a significant signal for a cysteine-rich region (N-score = 9.186, E-value = 0.014). Further, we utilized string-db.org to identify known connections and perform gene ontology of the 12 overlapping proteins. These proteins are strong interactors with each other, as shown in the protein interaction network (**Fig. 4C** and **Supplementary spreadsheet 1**). We reported the enrichment p-value for the number of interactions and ontology terms with an FDR of 5% or lower. Thus, these 12 SASP proteins are significantly regulated by ADAM19 and potentially targeted through a common cysteine-rich binding motif. In summary, in this experiment, we have identified ADAM19-altered secretory factors in senescence cells. Based on the sequence similarity of ADAM19’s cleavage site using motif analysis, we identified 12 SASP proteins as potentially directly targeted by ADAM19 knockdown.

**Figure 4.**
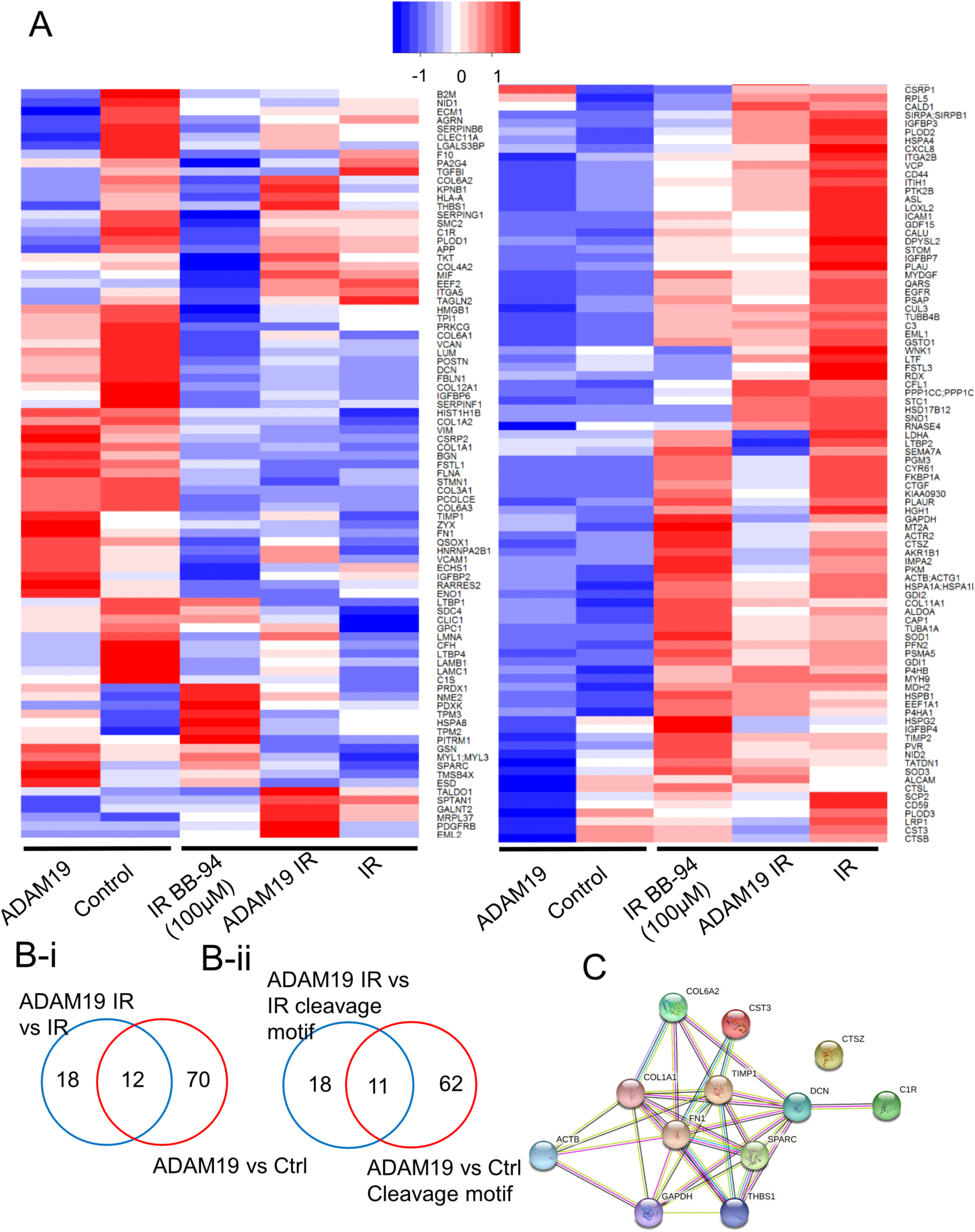
ADAM19 knockdown or inhibition alters secretory factors from senescent cells. **(A)** Heat map showing changes in the secretome composition from senescent IMR-90 cells with or without ADAM19 shRNA. **(B-i)** Venn diagram shows altered protein between ADAM 19 vs IR and IR only groups. **(B-ii)** Venn diagram shows altered protein with cleavage motif in groups between ADAM 19 vs IR and ADAM19 vs control. **(C)** Protein network 12 commonly altered protein in groups between ADAM 19 vs IR and IR.

## Discussion

Unresolved DNA damage is a known aging driver, causing a progressive decline in organismal and tissue function. Studying DNA damage in cell cultures or rodents has limitations, as cell cultures cannot recapitulate complex tissue microenvironments, and rodents are expensive and time-consuming. We recently demonstrated the fruit fly *Drosophila melanogaster* as an expedient model for studying genotoxic stress and its impact on tissue function and longevity[16]. We demonstrated its viability in studying radiation damage to the intestine, leading to increased intestinal permeability and reduced lifespan.

One response to genotoxic stressors such as radiation is cellular senescence, a state of permanent mitotic arrest characterized by the secretion of many proteins, including pro-inflammatory ones. Here, we report that in addition to increased intestinal permeability, exposure to ionizing radiation also led to an increase in persistent DNA damage foci in the gut of adult flies. We also observed an increase in the β-galactosidase expression in the gut of flies exposed to 100Gy ionizing radiation[16]. This increase in the expression of two well-characterized senescence markers was also observed in twenty-one-day-old flies, suggesting an age-dependent accumulation of senescent cells in the fly gut. These data correlate well with the previously reported increase in intestinal permeability of irradiated and old flies, suggesting a link between intestinal homeostasis and senescence burden [16][31]. Previously, simultaneous activation of the Ras oncogene and mitochondrial dysfunction in Drosophila imaginal epithelium has been shown to induce cell cycle arrest and secretion of proinflammatory SASP [55]. Our data supports this and further confirms the involvement of cellular senescence in flies as a physiological response to stress and tissue dysfunction caused by it in response to radiation-induced damage and aging. The practicality of the fly models, their amenability to genetic manipulation, and the viability of quantifiable assays like intestinal permeability offer opportunities to study cellular senescence in flies. In addition to senescence, another physiological response is apoptosis, in our previous manuscript, we reported an increase in the apoptotic clearance of cells with DNA damage at least up to day three after irradiation and a significant decrease in gut size 14 days after irradiation, our results in this manuscript show that the at least a subset of cells in the gut express several makers of senescence.

We identified naturally occurring gene variants that modulate intestinal damage upon irradiation by GWAS study. One gene revealed in our analysis was the highly conserved ADAM19/Meltrin β. <u>A D</u>isintegrin <u>and M</u>etalloproteases (ADAMs) are highly conserved in several species [19] [18], with their primary function being ectodomain shedding. Members of this class have a wide range of regulatory functions. They can proteolytically release several cytokines and growth factors (TNF-α, TGF-β), receptors (IL-1R-II), adhesion molecules, and enzymes from the plasma membrane [20–23,25,56]. Our results showed that tissue-specific knockdown of *meltrin β*/ADAM19 using an enterocyte-specific drug-inducible gene switch driver (5966 GS-Gal4) in adult flies decreases gut permeability, as determined by the Smurf assay. The increased expression of ADAM19 in the biopsy of patients with ulcerative Colitis and Crohn’s disease suggests its role in inflammatory pathologies causing leaky intestines in humans[36]. In this context, the heightened risk of leaky gut diseases like enteritis and proctitis in cancer therapy patients has been well documented[57,58]. Our results did confirm increased intestinal permeability in mice following a single treatment with doxorubicin[59]. We also observed an increase in the expression of proinflammatory SASP factors like *il6* and *tnfa* and an increase in the expression of p21 in the colon in doxorubicin-treated mice. This data aligns with previous reports showing increased expression of multiple senescent markers, including p21, p16, Tp53, and β-galactosidase, in the colon of aged mice compared to young mice[60].

ADAM19 knockout mice have cardiac defects and die soon after birth [61], so to investigate if its effects on intestinal permeability are conserved in mammals, we used pharmacological inhibition of ADAM19 using BB-94. ADAM19 is sensitive to BB94 but not to tissue inhibitors of metalloproteases (TIMPs)[62]. Thus, we used BB94 to inhibit ADAM19 expression in mice models. Batimastat or BB-94 treatment has been shown to reduce inflammation in the rat model of colitis [63]. Additionally, intraperitoneal injection of BB94 has been shown to minimize/restore blood-brain permeability disrupted in mice treated with TNF-α[64]. However, BB94 can function as the broad-spectrum metalloprotease inhibitor, and the possibility of its effect on intestinal permeability might be due to its impact beyond inhibiting just ADAM19 cannot be ruled out.

Nonetheless, taken together, our data offers ADAM19 as a viable target for several pathologies where disruption in gut homeostasis is involved, but it also provides metalloprotease inhibition as a viable therapeutic intervention.

Interestingly, ADAM19/*meltrin β* knockdown (or inhibition with BB-94) in human fetal lung fibroblasts IMR-90 also reduced the percentage of cells expressing β galactosidase. Interestingly, these interventions also decreased the expression of prototypical SASP factors like IL-6, IL-8, and IL-1β. In contrast, it did not affect the persistent DNA damage foci, DAMP maker HMGB1 or leminB1, thus indicating the role of ADAM19/melting β in regulating pro-inflammatory SASP from senescent cells. Our proteomics analysis of the conditioned media collected from senescent cells further supports the broad effect of ADAM19 inhibition or its knockdown on several SASP factors. As stated earlier, ADAM19 is a membrane-bound metalloprotease involved in shedding. Its role in regulating SASP release from senescent cells is conceivable. Its close homolog ADAM17 has also been shown to regulate SASP expression from senescent cells[65]. Morancho et al. concluded that senescent cells might tightly regulate ectodomain shedding. Thus, ADAMs like ADAM19 might play a vital role in regulating pro-inflammatory SASP in senescent cells. Our data shows that ADAMs like ADAM19 inhibitors might be a novel class of senomorphics.

Growing evidence suggests that gut permeability can be detected by measuring specific plasma components [66,67]. In our experiment, the knockdown of ADAM19 was shown to protect against increased gut permeability and reduce inflammatory signature. Additionally, we identified 29 senescence-associated secretory proteins (SASPs) in the culture media of IMR-90 cells following ADAM19 knockdown. Of these, 19 have been previously reported as SASPs [13,14]. Notably, GDF15 (Growth Differentiation Factor 15), TIMP1(Tissue Inhibitor of Metalloproteinases-1), and ICAM1(Intercellular Adhesion Molecule 1) were among the identified SASPs, all of which are known regulators of inflammation [68–70]. In a murine model of Crohn’s disease, TIMP1 knockdown has been shown to attenuate gut inflammation [71]. Moreover, a cell line-based study highlighted a potential role for TIMP1 in maintaining tight junctions and thus preserving gut integrity [72]. ICAM1 has been identified as a risk factor for inflammatory bowel disease (IBD) [73]. UPAR protein is another SASP from our study reported as a potential target to improve the intestinal epithelial barrier in IBD[74]. We also detected a few collagens (CO6A1, CO6A2, CO1A1, CO1A2), which are a key component in the extracellular matrix and known as important factors for wound healing processes in response to tissue injury[75]. Other proteins like TSP1 (Thrombospondin-1) and LTBP2 (Latent Transforming Growth Factor Beta Binding Protein 2) are involved in tissue remodeling and fibrosis, which are processes linked to gut inflammation and the chronic tissue damage seen in IBD[76–78]. These SASPs could be key players in gut diseases through their roles in inflammation, fibrosis, immune responses, and potential risk factors for gut integrity.

Our data shows that ADAM19 may be important in mitigating the inflammatory response to intestinal damage and cellular senescence. We also reported the role of batimastat as a senomorphic drug. However, whether it does so by inhibiting only ADAM19 or other metalloproteases remains to be explored. Identifying several SASP factors that may be involved in the pro-inflammatory response in gut permeability or senescence may shed light in future research to explore the precise role of ADAM19 in gut health and use it as a potential biomarker for gut integrity and therapeutic target for gut diseases.

## Methods

### Fly culture and stocks

Flies were reared on a standard laboratory diet (Cornmeal 8.6%, Yeast 1.6%, Sucrose 5%, Agar 0.46%, mixture of propionic acid and orthophosphoric acid 1%). Eclosed adults were transferred to standard food after 14 days of egg laying. For *Gene-Switch Gal4* drivers, 100μM of RU486 (dissolved in 95% ethanol) was used (the media was then referred to as ’+RU486’). The control diet contained the same volume of 95% ethanol and was referred to as ’-RU486’. The following fly strains were obtained from Bloomington *Drosophila* stock center W^1118^, *meltrin β* RNAi (58331), and 5961-GS, a kind gift from Dr. Heinrich Jasper.

### Radiation exposure

Adult, female, 5-day-old flies were exposed to X-rays at 320 kV and 10 mA for 10 min to achieve 100 Gy and maintained on eighter + RU486 and - RU486 diets.

### Smurf gut permeability assay

We performed the smurf assay as previously described with slight modifications[31]. Briefly, we introduced 25 flies into an empty vial that held a 2.0 cm x 4.0 cm piece of filter paper. To serve as their food source, we applied 300 μl of a 2.5% blue dye solution (FD&C #1) in a 5% sucrose solution to saturate the paper. After a 24-hour incubation at 25°C, we counted the number of flies displaying visible blue dye outside their intestines to identify as “Smurf” flies.

### Immunohistochemistry

Immunohistochemistry was performed post-14 days of X-ray irradiation. Briefly, flies were dissected for gut in PBS and immediately fixed with 4% PFA for 30 minutes. Samples were washed for 10 minutes three times with PBS (Phosphate buffer saline) and then incubated with PT (PBS, 0.3% triton X-100) for 30 minutes. Samples were washed for 10 minutes three times with PBS, and blocking was performed with 5% BSA in PT for 1 hour at room temperature. Samples were incubated with primary antibody overnight at 4°C, washed for 10 minutes three times with PT, and incubated with secondary antibody for 2 hours at room temperature and post-incubation wash three times with PT. Nuclei were stained using DAPI. Samples were mounted with Fluromount G (southern biotech), and images of midgut were captured in a Zeiss 780 confocal microscope. The following antibodies were used in this study: anti-GLB1 (abcam#203749: 1/200), anti-mouse γ-H2Av (DSHB: 1/200), anti-rabbit Alexa fluor 488 (Cell Signaling Technology: 1/500) and anti-mouse Alexa fluor 555 (Cell Signaling Technology: 1/500). These were analyzed using Image J number by counting γ-H2AX positive cells and total intensity of β galactoside within an ROI of the 11583-unit area and normalized with a total nucleus within this area. The nucleus of more than two foci was marked as γ-H2AX positive cells.

### Bioinformatics analysis

Pathway analysis of Senescence-Associated Secretory Phenotype (SASP) proteins identified through proteomics with and without Adam19 knockdown (KD) was performed using tools from reactome.org[79]. Significant proteins with an Adam19 cleavage site identified through proteomics in each group, Adam19 vs control and Adam19 with IR vs control with IR, were compared to lists of identified SASP proteins[13,14]. The SASP proteins of each group were then coded as 1, and non-SASP proteins as 0, and the groups were separately inputted into the Pathway Browser analysis tool, reactome.org, without including interactors.

### Cell Culture

IMR-90 fetal lung fibroblasts (ATCC, USA: Cat# CCL-186) cultured in IMR-90 cells were cultured in complete media containing Dulbecco’s modified Eagle’s medium (DMEM) (Corning; Cat# 10-013-CV) supplemented with 10% fetal bovine serum (FBS) (Millipore Sigma, US; Cat# F4135) and 1x penicillin-streptomycin (Corning; Cat# 30-001-CI). The cells were a 5% CO2 and 3% O2 incubator.

### Knockdown by ADAM19 using lentiviral shRNA

We selected MISSION shRNA Plasmid (SHCLNG-NM_023038, Sigma, USA) to stable knockdown ADAM19. The viral packaging and introduction of lentivirus into host cells were done according to the Addgene pLKO.1 protocol (Addgene, USA). The selection of cells stably expressed ADAM19 shRNA, and the control-shRNA started 72 h post-transfection. The growth medium was replaced with a fresh selection medium containing 10 μg/mL of puromycin. The puromycin-containing medium was refreshed every other day for four days. QRT-PCR was performed to determine the expression of ADAM19.

### Senescence induction

IMR-90 cells were treated with 300nM of doxorubicin Hydrochloride (Millipore Sigma, USA; Cat# 504042) in DMEM complete media for 24h, and cells were maintained in DMEM complete media by changing fresh media every other day for additional eight days, doxorubicin treated and non-senescent control cells were treated with low serum DMEM (0.2% FBS DMEM media) for one day before harvesting cells for various assays.

### Senescence-associated ß-galactosidase staining

Senescent-associated ß-galactosidase activity in IMR-90 cells was assessed using the Senescence Detection Kit (BioVision; Cat# K320), following the manufacturer’s instructions. The percentage of SA-ß-gal-positive cells was counted manually. Four fields were quantified per well (n = 3).

### Real-Time Quantitative PCR

Cell pellets of senescent IMR-90 fibroblasts were collected nine days after doxorubicin treatment of IMR-90 cells. The total RNA was isolated using Quick-RNA MiniPrep (Zymo Research; Cat# R1055), following the manufacturer’s protocol. cDNA was synthesized using the High-Capacity cDNA RT kit (Life Technologies). Transcripts were analyzed using a Roche LightCycler 480 II in 384-well plates and the UPL probe system. Bioline SensiFast Probe No-ROX was used as a master mix. Each gene’s mean cycle threshold (Ct) was normalized to tubulin levels in the same sample (delta Ct). Unpaired two-sample t-tests determined differences in mean delta Ct values between treatment groups. The delta-delta calculated the fold change. Ct method (fold = 2ddCt).

### Mouse model

C56BL/6 mice were purchased from The Jackson Laboratory and housed under controlled temperature (22-24° C), humidity (40-60%), and a 12-hour light-dark cycle at the Buck Institute. All animal procedures were approved by the Buck Institute Institutional Animal Care and Use Committee (IACUC). The six-month-old male mice received a single intraperitoneal (IP) doxorubicin injection (10 mg/kg) to induce widespread senescence. A sub-set of mice received BB-94 *i.p* injection (5 mg/Kg or 20 mg/Kg) once a week till the completion (6 weeks) starting 48 hours after Dox injection. After six weeks, the animals were euthanized by CO2 inhalation followed by cervical dislocation, and tissues were harvested and flash-frozen in liquid nitrogen.

### Albumin ELISA

The albumin concentration in freshly collected fecal pellets from mice was determined by a standard ELISA test using the manufacturer’s instructions (Bethyl Laboratories, # E99-134). Samples were collated from single-housed mice in cages without bedding for several hours to collect fecal pellets from individual mice. The fecal pellets were re-suspended in 1xPBS for ELISA.

### Collection of Secreted Soluble Proteins for Mass Spectrometry

Proteins secreted into a serum-free medium over a 24-hr period were collected. The conditioned medium was centrifuged at 10,000 x g at 4° C for 30 minutes to remove debris. The supernatant containing soluble secreted was saved and processed for mass spectrometry.

### Proteomic Sample Preparation

#### Chemicals

Acetonitrile (#AH015) and water (#AH365) were from Burdick & Jackson. Iodoacetamide (IAA, #I1149), dithiothreitol (DTT, #D9779), formic acid (FA, #94318-50ML-F), and triethylammonium bicarbonate buffer 1.0 M, pH 8.5 (#T7408) were from Sigma Aldrich, urea (#29700) was from Thermo Scientific, sequencing grade trypsin (#V5113) was from Promega and HLB Oasis SPE cartridges (#186003908) were from Waters.

#### Protein concentration and quantification

Samples were concentrated using Amicon Ultra-15 Centrifugal Filter Units with a three-kDa cutoff (MilliporeSigma #UFC900324) as per the manufacturer’s instructions and transferred into 8M urea/50 mM triethylammonium bicarbonate buffer at pH 8. Protein quantitation was performed using a BCA Protein Assay Kit (Pierce #23225).

#### Digestion

Aliquots of each sample containing 25-100 µg protein were brought to equal volumes with 50 mM triethylammonium bicarbonate buffer at pH 8. The mixtures were reduced with 20 mM DTT (37°C for 1 hour), then alkylated with 40 mM iodoacetamide (30 minutes at RT in the dark). Samples were diluted 10-fold with 50 mM triethylammonium bicarbonate buffer at pH eight and incubated overnight at 37°C with sequencing grade trypsin (Promega) at a 1:50 enzyme: substrate ratio (wt/wt). <u>Desalting</u>: Peptide supernatants were collected and desalted with Oasis HLB 30 mg Sorbent Cartridges (Waters #186003908, Milford, MA), concentrated, and re-suspended in a solution containing mass spectrometric ‘Hyper Reaction Monitoring’ retention time peptide standards (HRM, Biognosys #Kit-3003) and 0.2% formic acid in water.

### Mass Spectrometry Analysis

Samples were analyzed by reverse-phase HPLC-ESI-MS/MS using the Eksigent Ultra Plus nano-LC 2D HPLC system (Dublin, CA) combined with a cHiPLC system directly connected to an orthogonal quadrupole time-of-flight SCIEX TripleTOF 6600 or a TripleTOF 5600 mass spectrometer (SCIEX, Redwood City, CA). Typically, mass resolution in precursor scans was ∼ 45,000 (TripleTOF 6600), while fragment ion resolution was ∼15,000 in ‘high sensitivity’ product ion scan mode. After injection, peptide mixtures were transferred onto a C18 pre-column chip (200 µm x 6 mm ChromXP C18-CL chip, 3 µm, 300 Å, SCIEX) and washed at 2 µl/min for 10 min with the loading solvent (H2O/0.1% formic acid) for desalting. Peptides were transferred to the 75 µm x 15 cm ChromXP C18-CL chip, 3 µm, 300 Å, (SCIEX), and eluted at 300 nL/min with a three hours gradient using aqueous and acetonitrile solvent buffers.

All samples were analyzed using data-independent acquisitions (DIA), specifically variable window DIA acquisitions[53,80,81]. In these DIA acquisitions, windows of variable width (5 to 90 m/z) are passed in incremental steps over the full mass range (m/z 400-1250). The cycle time of 3.2 sec includes a 250 msec precursor ion scan followed by 45 msec accumulation time for each of the 64 DIA segments. The variable windows were determined according to the complexity of the typical MS1 ion current observed within a certain m/z range using a SCIEX ‘variable window calculator’ algorithm (more narrow windows were chosen in ‘busy’ m/z ranges, wide windows in m/z ranges with few eluting precursor ions). DIA tandem mass spectra produce complex MS/MS spectra, a composite of all the analytes within each selected Q1 m/z window. All collected data was processed in Spectronaut using a pan-human library that provides quantitative DIA assays for ∼10,000 human proteins.

### Processing, Quantification, and Statistical Analysis of MS Data

DIA acquisitions were quantitatively processed using the proprietary Spectronaut v12 (12.020491.3.1543) software from Biognosys. A pan-human spectral library was used for Spectronaut processing of the DIA data. Quantitative DIA MS2 data analysis was based on extracted ion chromatograms (XICs) of 6-10 of the most abundant fragment ions in the identified spectra. Relative quantification was performed by comparing different conditions (senescent versus control) to assess fold changes. To account for variation in cell number between experimental groups and biological replicates, quantitative analysis of each sample was normalized to cell number by applying a correction factor in the Spectronaut settings. There were three replicates for each experimental condition: X-irradiated senescent fibroblasts, mock-irradiated non-senescent fibroblasts, Adam19 shRNA-treated senescent fibroblasts, BB94-treated senescent fibroblasts, Adam19 shRNA-treated non-senescent fibroblasts. Significance was assessed using FDR-corrected q-values<0.05.

### Data Visualization

Heatmaps were visualized in R using the heatmap2 function of the ‘gplots’ package[82].

## Data availability

All raw files are uploaded to the Center for Computational Mass Spectrometry, MassIVE, and can be downloaded using the following link https://massive.ucsd.edu/ProteoSAFe/dataset.jsp?task=a8341a1bcba442e7ae151e15919ac92b (MassIVE ID number: MSV000085224). Data uploads include the protein identification and quantification details, spectral library, and FASTA file used for mass spectrometric analysis. MassIVE: MSV000085224 (https://massive.ucsd.edu/ProteoSAFe/dataset.jsp?task=a8341a1bcba442e7ae151e15919ac92b)

## Author contributions

Conceptualization (AS, PK, JC), methodology (SB, TAUH, BP, JALD, NB, JB, MW, BS, AS), investigation (SB, AS, TAUH, BP, JAL, NB, JB, MW), visualization (SB, AS), funding acquisition (SB, PK), project administration (AS, PK), supervision (AS, PK), writing – original draft (SB, AS, PK), review & editing: (SB, AS TAUH, PK)

## Figure legends

**Supplementary Figure 1:**
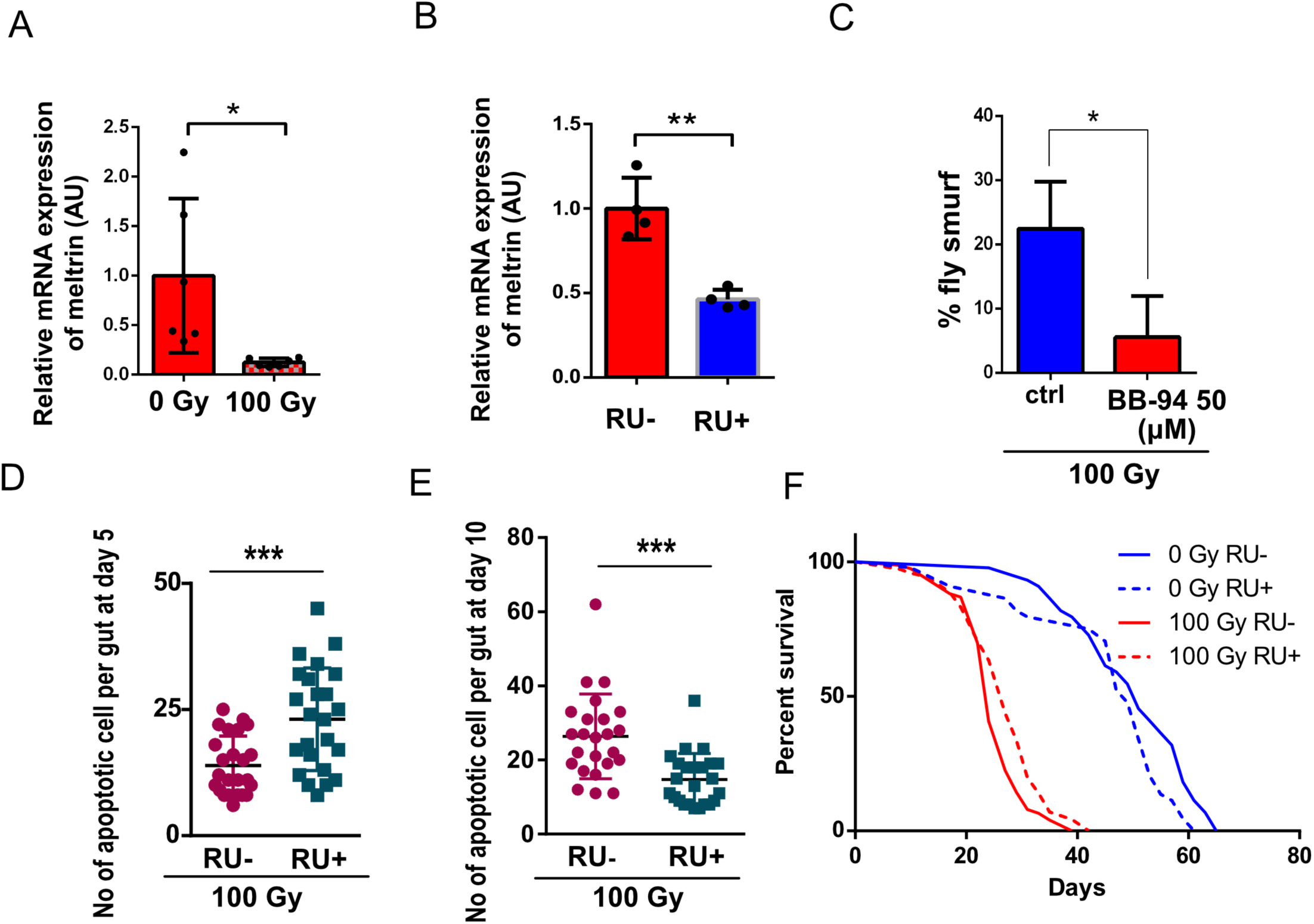
**(A)** Relative mRNA expression of meltrin detected by qRT PCR in flies upon X-Ray irradiation. **(B)** Relative mRNA expression of meltrin detected by qRT PCR in the gut of meltrin RNAi flies. **(C)** % of Smurf-positive flies with leaky gut upon 50 µM BB-94 treatment after 100 Gy X-Ray exposure. **(D)** Quantification of apoptotic cells upon meltrin knockdown and 5-day post-X-ray irradiation. **(E)** Quantification of apoptotic cells upon meltrin knockdown and 10-day post-X-ray irradiation. **(F)** The lifespan of *Drosophila* in flies with X-ray exposure (100 Gy), No X-ray exposure (0 Gy), meltrin knockdown (RU+) in ECs, and no meltrin knockdown control (RU-). Error bars indicate SEM. The student’s t-test determined the significance level by asterisks: * for p < 0.05, ** for p < 0.01, *** for p < 0.001, and **** for p < 0.0001.

**Supplementary Figure 2:**
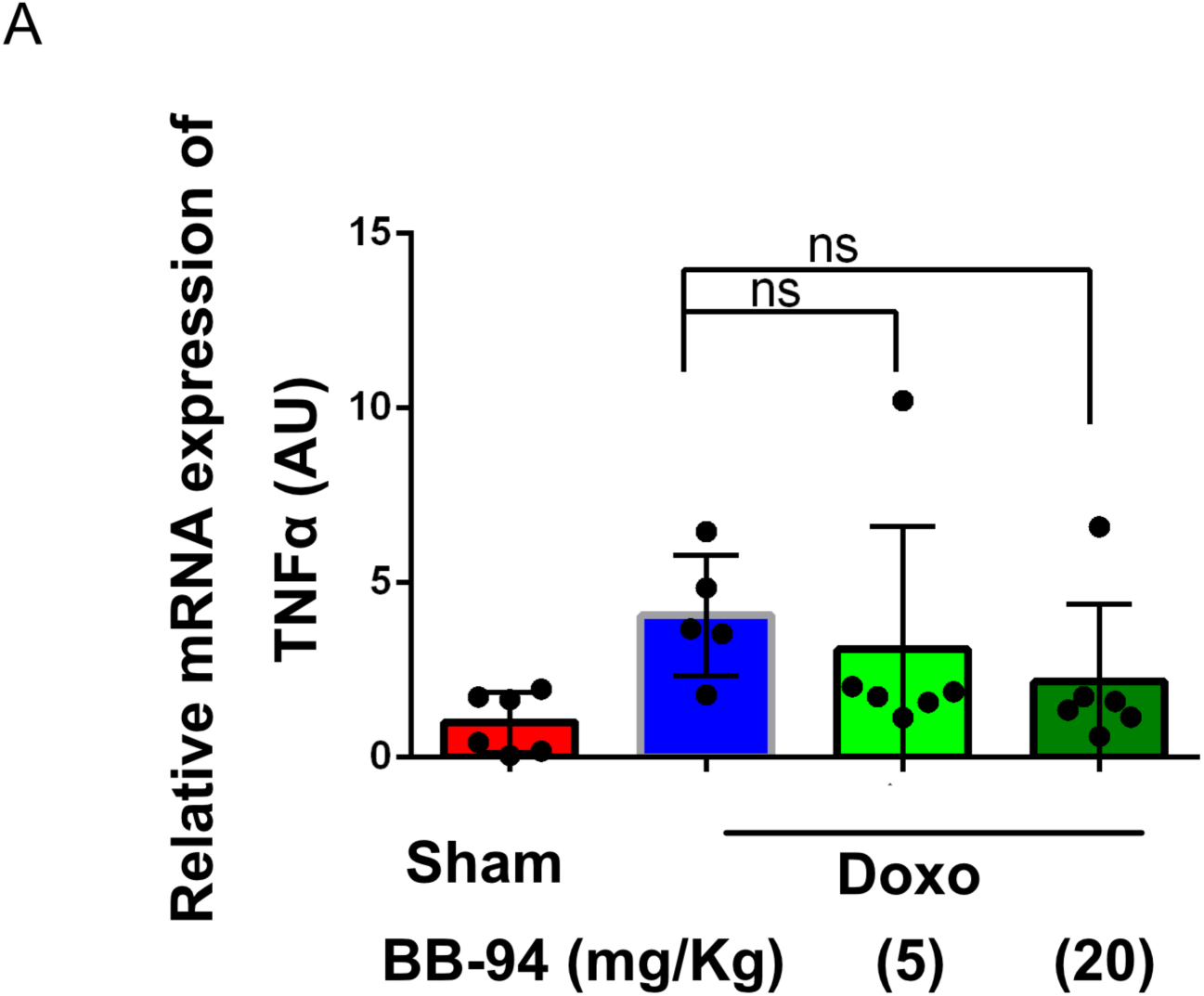
**(G)** Relative mRNA expression of TNFα in mouse colon tissues. Error bars indicate SEM. The significance level was determined by ANOVA and denoted by ns (no significance)

**Supplementary Figure 3:**
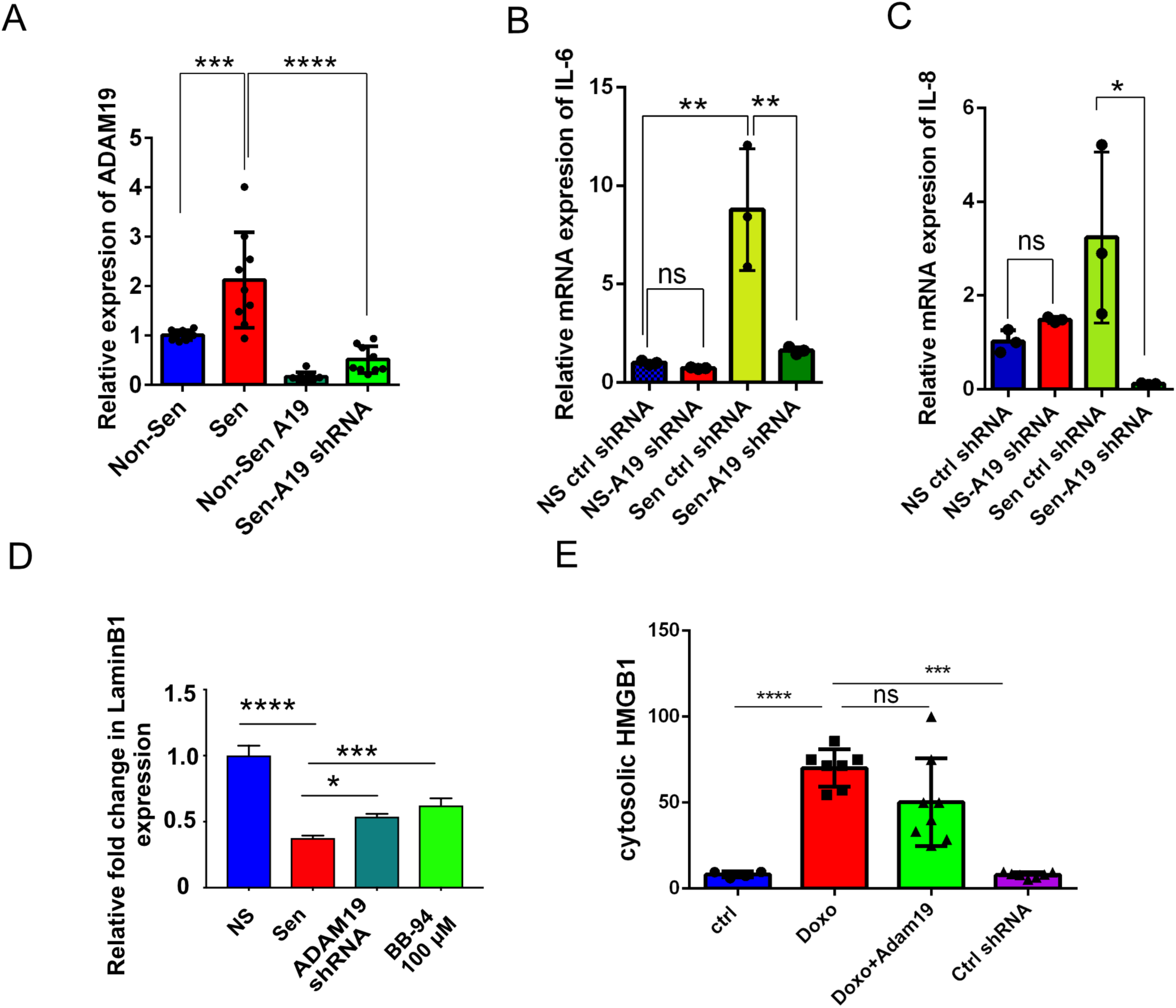
**(A)** ADAM19 expression upon senescence and non-senescence with ADAM 19 0r shRNA control treatments. **(B)** Relative mRNA expression of IL-6 in IMR90 cells upon senescence and non-senescence with ctrl shRNA and ADAM19 shRNA transduction. (C) Relative mRNA expression of IL-8 in IMR90 cells upon senescence and non-senescence with ctrl shRNA and ADAM19 shRNA transduction. (D) Lamin B expression in control, senescence, senescence with ADAM19 shRNA expressing and with BB-94 treatment cells. (E) % cells count with cytosolic HMGB1 in control, senescence, and senescence with ADAM19 shRNA expressing cells. Error bars indicate SEM. The significance level was determined by ANOVA and denoted by asterisks: * for p < 0.05, ** for p < 0.01, *** for p < 0.001, and **** for p < 0.0001. In A, B, C and D “NS” represent non senescence and ‘Sen’ represent senescence.

